# Comparison of the 4-helix bundle domains of perilipin 3 and perilipin 4 identifies features that contribute to lipid droplet binding

**DOI:** 10.64898/2026.08.15.745046

**Authors:** Cyril Moulin, Bayane Sabbagh, Amel Bahloul, Nicolas Fuggetta, Romain Gautier, Alenka Čopič

## Abstract

The perilipins generally represent the most abundant lipid droplet (LD) surface proteins in mammalian cells and can target LD subpopulations within the same cell. They are characterized by a conserved organization of disordered and folded regions, as well as a number of divergent features, which contribute to differences in perilipin function and LD targeting. Here, we focus on the C-terminal 4-helix bundle (4HB) domain that is present in all perilipins except for PLIN1. Using biochemical and *in silico* approaches, we show that the 4HB of PLIN3 is a stably folded domain and interacts with lipid surfaces *in vitro* and with LDs in model cells. The αβ-subdomain at the bottom of the helical bundle is required for the binding to LDs, but not for the 4HB stability, suggesting that this region may promote direct interaction with the LD surface. In agreement, the 4HB of PLIN4, which does not contain an αβ- subdomain, does not bind to LDs. Overall, our work shows that small differences in perilipin structural features impact their differential targeting to LDs.

## Introduction

Perilipins are evolutionarily-conserved proteins that localize to the surface of lipid droplets (LDs) and often represent the most abundant LD surface proteins. They can regulate lipid hydrolysis in LDs via lipolysis, lipophagy or fatty acid trafficking, LD biogenesis, interaction with other organelles, LD stability or fusion. However, despite decades of research on perilipins, their exact functions and mode of interaction with LDs remain only partially understood. In mammals, the perilipin family is comprised of five proteins, PLIN1-PLIN5, which vary in terms of their tissue distribution and transcriptional regulation. PLIN1 and PLIN4 are highly expressed in mature adipocytes, where PLIN1 acts as a master regulator of lipolysis. PLIN5 fulfills a similar role in oxidative tissues. PLIN2 is highly expressed in liver and muscle tissue, whereas PLIN3 displays the most ubiquitous and stable expression profile (1–3).

The phylogenetic relationship between the perilipins has been well described (1, 4, 5). They can be defined by a common sequence organization: an N-terminal region, termed the PAT- domain, followed by a central repetitive region and a C-terminal 4-helix bundle (4HB) domain (6–8) (Fig. 1A). However, none of these features is strictly conserved among the mammalian perilipins (3). The PAT-domain is present in all perilipins except PLIN4, the repetitive region varies in length from 60 to 100 amino acids in all perilipins except PLIN4, where this region is extended to close to 1000 amino acids (9–13). Finally, the C-termini after the 4HB can be extended to include additional functional motifs, in particular in PLIN1 and PLIN5 (Fig. 1A) (14–17).

**Figure 1.**
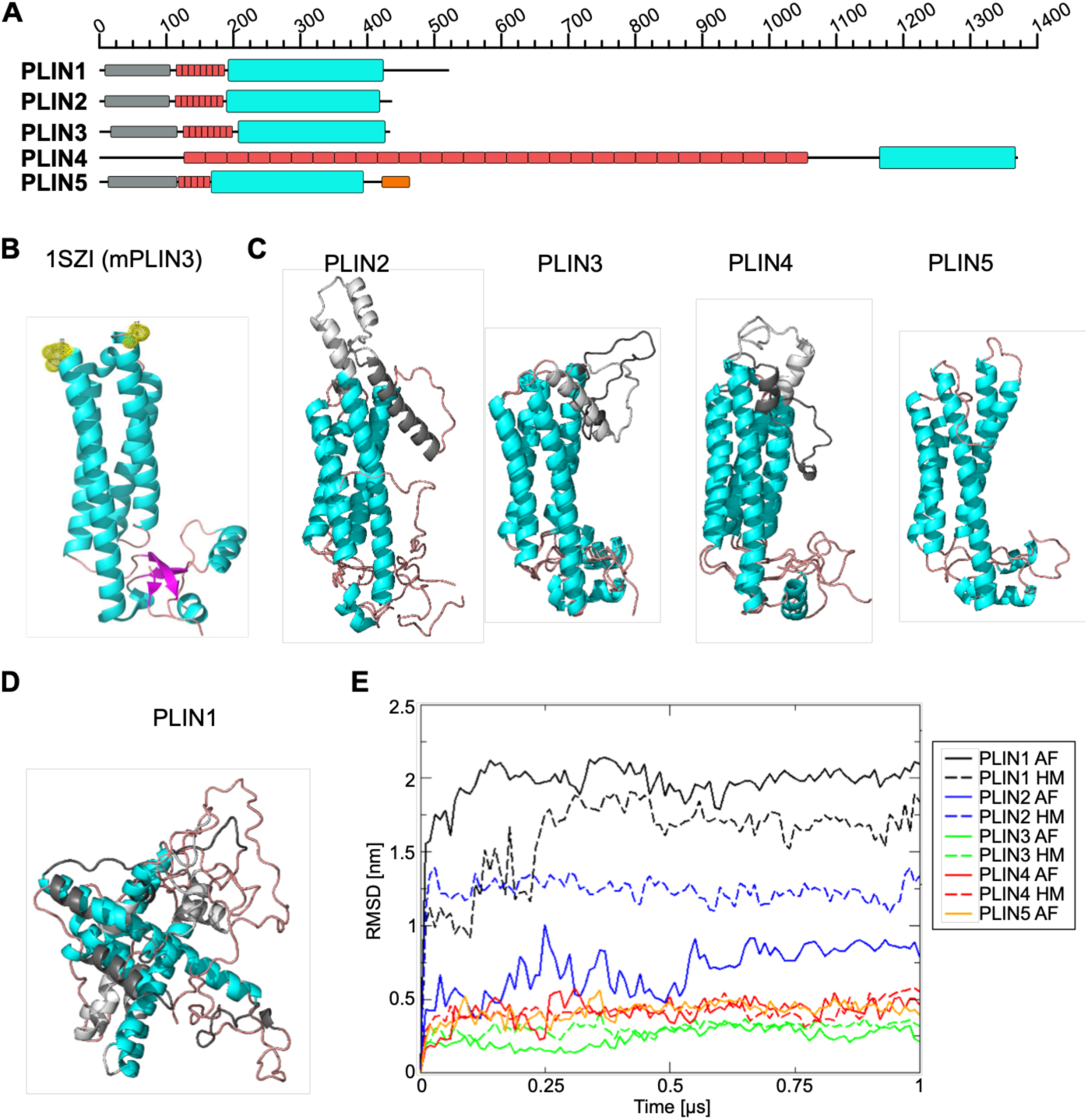
Comparison of 4HB regions in human perilipins. **(A)** Diagrams of human perilipins, showing the positions of the regions (in cyan) homologous to the 4HB domain from mouse PLIN3 (hPLIN1: L191-E413, hPLIN2: L191-E411, hPLIN3: N202-E427, hPLIN4: M1166-Q1371, hPLIN5: F165- P388). The repetitive AH regions are shown in red, N-terminal PAT domain in grey, and the C-terminal mitochondria-interacting region in PLIN5 in orange. **(B)** Crystal structure of the mouse PLIN3 4HB (I206- 431 PDB 1SZ) (33). Yellow spheres mark the missing part in the crystal structure (residues 289-320). **(C)** Structural models of 4HB regions of human PLIN2, PLIN3, PLIN4 and PLIN5 and PLIN2, obtained with AlphaFold or by homology modeling, after 1-μs equilibrations by MD simulations in solution. The two models are superposed, with the divergent regions shown in light (AlphaFold) or dark grey (homology modeling). **(D)** Superposition of the AlphaFold (grey or cyan) and homology (grey or cyan) models of human PLIN1 4HB region after 1-μs equilibration by MD simulations in solution. **(E)** The stability of 4HB structures from PLIN1-PLIN5 (AF and homology models) in solution was analyzed using all-atom MD simulations (supplemental movies 1-10). The root mean square deviation (RMSD) of the domain structure compared to the initial model is plotted over time.

Importantly, even within the same cell, different perilipins often do not target the same LDs. This was first observed in cultured adipocytes, where PLIN1 localizes to large central LDs, whereas PLIN2, PLIN3 and PLIN4 can be observed on smaller and more peripheral LDs (18, 19). In accordance, they do not interact with LDs in the same manner: PLIN3 and PLIN4 bind to LDs from the cytosol in response to LD growth after fatty acid feeding, and require the repetitive region, which folds into an amphipathic helix (AH) on the LD surface, for LD binding (11, 20–22). PLIN3 is also sensitive to diacylglycerol, which may promote its binding to the ER during LD-biogenesis via the N-terminal PAT domain (8, 23–25). By contrast, PLIN1 and PLIN2 stably associate with LDs, with PLIN1 relocating to the LD surface from the ER utilizing hydrophobic regions within its C-terminal part. Furthermore, perilipins show different preferences for LD core or lipid surface composition, which may be due to their mode and depth of insertion into the LD phospholipid monolayer (22, 26, 27).

To better understand how perilipins interact with LDs to regulate their function, it is essential to dissect the contribution of each of their structural features to LD binding. One region that is well conserved in most perilipins, except for PLIN1, is the C-terminal 4HB domain (Fig. 1A); however, there is conflicting evidence to what extent this domain contributes to LD binding (8, 28–32). The Hurley group determined the crystal structure of the 4HB domain from mouse PLIN3 in 2003, showing that it folds into a stable domain comprising four extended helices that form the bundle, and an α/β subdomain at the bottom of the bundle, composed of sequences upstream and downstream of the helical bundle (33). The 4HB motif is a structural feature that is present in many proteins or protein complexes (34). Hickenbottom et al. noted a strong resemblance between the 4HB of PLIN3 and some other proteins including α-catenin, cytochrome-c and apolipoprotein E, despite a lack of any sequence similarity. In apolipoproteins, the helical bundle can undergo large structural rearrangements when in contact with lipids, leading to the helices splaying apart to laterally interact with the lipid core of the lipoprotein particle (35). It was later proposed that a similar mechanism might operate in perilipins, with the α/β subdomain, which is not present in apolipoproteins, functioning as a zipper to stabilize the helical bundle (31); however, this mechanism was not experimentally demonstrated. Furthermore, purified 4HB of PLIN2 was shown to be structurally stable by its resistance to proteolysis or urea treatment, and could directly interact with charged liposome membranes or the plasma membrane in cells (28). The PLIN3-4HB could interact with phospholipid-covered oil droplets in a drop tensiometry assay, or with highly hydrophobic and charged liposomes in vitro, suggesting a synergistic effect with the N-terminal portion of PLIN3 (8, 24, 36); these results contrasted with the previously reported lack of binding of the C-terminus of PLIN3 to LDs (21, 31).

Another feature of the 4HB of PLIN3 is a hydrophobic cleft surrounded by residues within the bundle as well as some residues within the C-terminal region of the α/β subdomain, which are conserved in PLIN2, PLIN4 and PLIN5 (33). It was shown that PLIN2-4HB and PLIN5-4HB can bind fatty acids using the hydrophobic cleft, with PLIN5 mediating fatty acid transfer to the nucleus to promote signaling (29, 37). It has not been tested whether this might apply to all perilipins; however, this mode of function does not seem compatible with opening up of the bundle upon LD interaction.

Here, we first systematically analyze the folding and stability of the 4HB-homology regions in human perilipins using modeling and molecular dynamics (MD) simulations. With the exception of PLIN1, all perilipins (PLIN2, 3, 4 and 5) possess a 4HB bundle that remains stable in solution during 1-2 µs all-atom MD simulation. Previous work has shown that PLIN4 displays the most similar behavior to PLIN3 in cells, existing in a soluble cytoplasmic pool and relocalizing to LDs early during their biogenesis (18, 20, 22). However, we show that the 4HB of PLIN3 binds to LDs in cells and to LD-mimetic liposomes *in vitro*, whereas the PLIN4-4HB remains soluble under all conditions. This difference can be explained by the absence of a structured αβ-subdomain in the PLIN4-4HB; deletion of this subdomain in PLIN3-4HB renders it unable to bind to LDs, but does not change its overall stability. We therefore suggest that the αβ-subdomain, rather than stabilizing the perilipin 4HB structure, promotes the interaction with the LD surface.

## Results

### Comparison of 4HB regions of human perilipins

We first constructed models of the C-terminal (4HB) homology regions from the human perilipins PLIN1-PLIN5 (Fig. 1A and S1A) using AlphaFold or homology modeling based on the structure of mouse PLIN3-4HB (Fig. 1B) (33). The structural models were equilibrated during 1 µs all-atom MD simulations in solution using the Charmm36 force-field. The MD simulations of the 4HB-homology regions of PLIN2, PLIN3, PLIN4 and PLIN5, modeled by AlphaFold or by homology modeling, show that they all form stable structures composed of 4 helices that interact with each other along their lengths (Fig. 1C). The results obtained with the two models showed high level of agreement. In case of PLIN2, PLIN3 and PLIN4, some differences were observed at the top part of the bundle, most strongly in the case of PLIN2, where helix 1 is either extended or split into another helix connected by flexible loops. Compared to the experimentally determined structure, an additional helix (here termed helix 5) is present in PLIN3, which is not part of the bundle and is connected with flexible loops; this entire region could not be resolved in the crystal structure of the mouse PLIN3-4HB (33). Differences in the structural models from PLIN2 to PLIN5 can also be observed at the base of the helical bundle, which contains additional short α-helices and β-sheets in the PLIN3 4HB structure, contributed by sequences upstream and downstream of the helical bundle region. This region was named the αβ-subdomain by Hickenbottom et al. (33), and will be discussed further later in the manuscript.

By contrast, the corresponding region in PLIN1 is more divergent in sequence and does not form a stable helical bundle structure; it cannot be superimposed with the 4HB from PLIN3 (Figs. 1D and S1B). This result is in agreement with analyses showing the presence of more hydrophobic segments in the PLIN1 4HB region, which are important for PLIN1 LD targeting via the ERTOLD pathway (32, 38, 39). The PLIN1 4HB may thus not be able to stably fold in solution. Indeed, purification of full-length PLIN1 from *Escherichia coli* required the addition of urea as the protein was not soluble in water-based buffers, likely due to its higher hydrophobicity (22).

The MD simulations of the 4HB-homology structures from different perilipins in solution are presented in supplementary data (Video S1-S9). We analyzed the room mean square deviation (RMSD) of the two models for each perilipin over time, compared to the initial model (only AlphaFold model has been simulated for PLIN5). The RMSD of the 4HB structural models of PLIN3, PLIN4 and PLIN5 showed minor changes over the course of the simulation. A larger change was observed for PLIN2, which also showed a larger difference between the AlphaFold and homology-based models. Finally, the predicted structure of the corresponding region of PLIN1 showed a much larger fluctuation over time, suggesting that this sequence does not form a stably folded structure in solution (Fig. 1E).

### The 4HB of PLIN4 does not bind to liposomes, contrary to PLIN3-4HB

PLIN3 and PLIN4 display similar behaviors in cells, existing as stable cytosolic proteins and translocating to the LD surface upon induction of LD formation (20, 18, 21, 22). Their ability to interact with the LD surface has been shown to depend on their repetitive central regions and the N-terminal PAT domain in the case of PLIN3 (8, 11, 12, 21, 31); however, some studies suggest that the 4HB of PLIN3 may also contribute to LD binding (8, 36). Following the prediction that the C-terminal region of PLIN4 folds into a 4HB similar to that of PLIN3 4HB, we wanted to further compare the properties of these two domains. We expressed and purified PLIN3 4HB and PLIN4 4HB from *E. coli* as glutathione S-transferase (GST) fusion proteins, and purified them using a three-step purification protocol (Fig. S2). After GST-affinity chromatography, the proteins were eluted by proteolytic cleavage between the tag and the protein using a highly specific TEV cleavage site. In the final step, the non-tagged proteins were further purified by size-exclusion chromatography, which yielded highly-pure final proteins that eluted as monomers and also as oligomers, in agreement with previous studies on PLIN3-4HB (33). Protein peaks corresponding to monomeric proteins were used in subsequent experiments, and the folded state of the purified proteins was verified by circular dichroism (CD) (Fig. S2E, F).

We used a sucrose flotation assay to assess the ability of purified proteins, PLIN3-4HB, full length PLIN3 (22) and PLIN4-4HB, to directly bind to liposomes (Fig. 2A-C). We used liposomes composed of diphytanoyl (diphy) phospholipids and a high amount of negative charge, up to 50% phosphatidylserine (PS), to promote maximum protein binding (8, 11). The branched methyl groups of diphy phospholipids prevent tight lipid packing, thereby mimicking an LD-like surface (40). In some experiments, we also use dioleoyl (DO) liposomes for comparison. In the absence of liposomes, proteins remained in the bottom (B) fraction (Fig. 2A,D), whereas proteins bound to liposomes were recovered in the top (T) fraction (Fig. 2E-F). As previously shown, both full-length PLIN3 and PLIN3-4HB showed binding to diphy liposomes containing 50% or 10% PS, with more binding observed with the full-length protein (Fig. 2A,B,E). By contrast, PLIN4-4HB did not show any binding to these liposomes, because it remained in the bottom fraction (Fig. 2C, E). PLIN4-4HB also did not bind to liposomes composed of dioleoyl phospholipids (DOPC, DOPS), whereas PLIN3-4HB exhibited some binding, as previously shown (Fig. 2F) (8).

**Figure 2.**
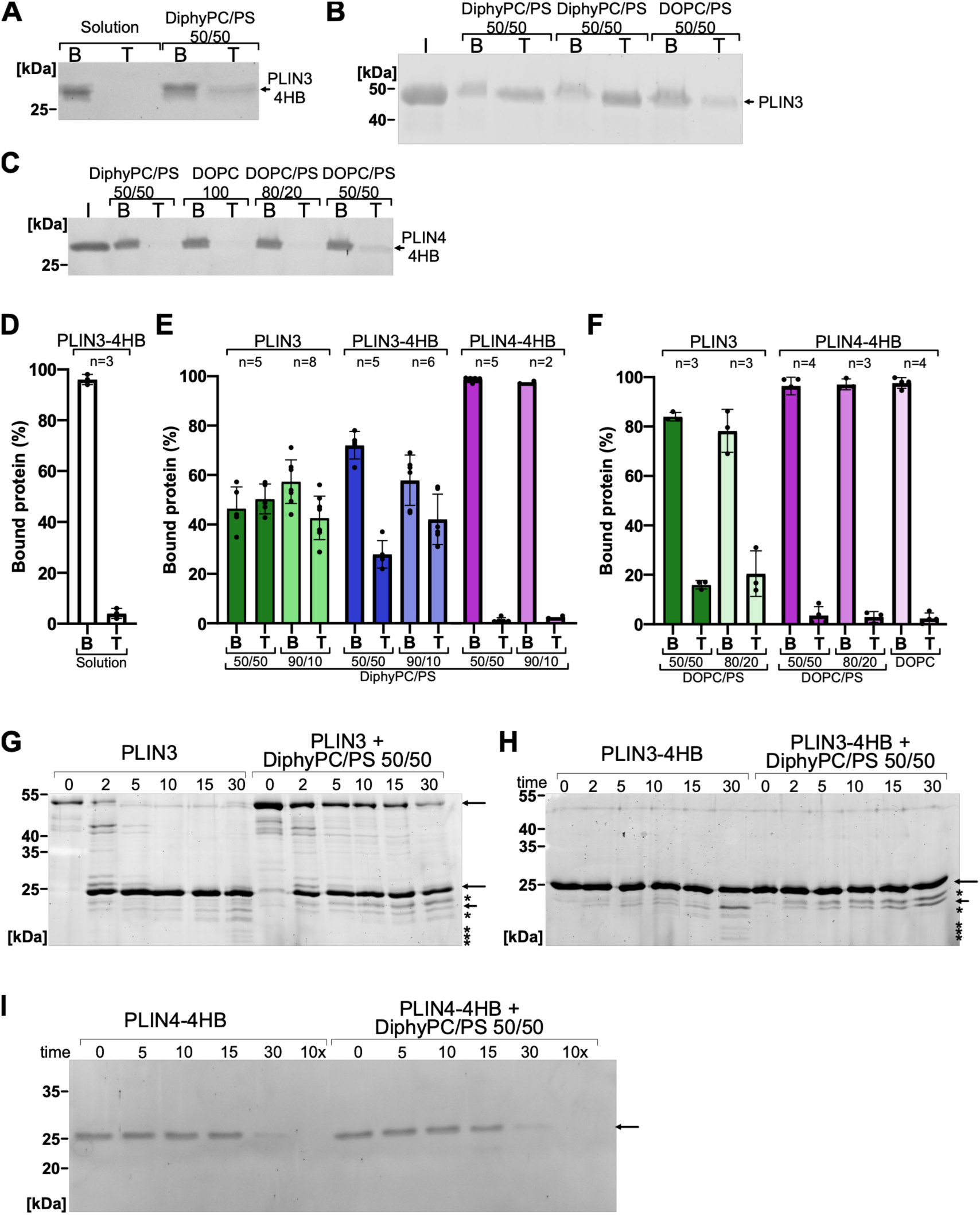
Comparison of liposome binding and stability of human PLIN3 4HB and PLIN4 4HB. (A-C) Purified PLIN3-4HB (A), PLIN3(full length) (B) and PLIN4-4HB (C) (2.2 - 3.5 μM) were incubated with liposomes ([lipid] = 500 μM) containing diphytanoyl (Diphy) or dioleoyl (DO) phospholipids at the indicated ratio of PC and PS headgroups, followed by flotation on sucrose gradients. Input (I), bottom (B) and top (T) fractions were analyzed by SDS-PAGE and proteins were stained with Sypro Orange (B). In control experiments without liposomes (A, first two lanes), all proteins remain at the bottom. **(D-F)** Quantification of liposome binding experiments as shown in A-C. Data are presented as mean ± SD, with data points representing individual experiments. The number of replicates and the liposome composition for each experiment is indicated above and below the bar graphs, respectively. **(G-I)** The stability of PLIN3 (G), PLIN3-4HB (H) and PLIN4-4HB (I) in solution or in the presence of Diphy liposomes (50% PC, 50% PS) was analyzed by limited proteolysis with trypsin (1 μg/ml). Samples were retrieved at indicated time points: 0, 2, 5, 10, 15 and 30 min for PLIN3 and PLIN3-4HB, and 0, 5, 10 and 30 min for PLIN4-4HB; a 10x (10 μg/ml) trypsin reaction for 30 min was included for PLIN4-4HB. Samples were analyzed by SDS-PAGE and stained with SyproOrange. Arrows mark protein bands that are common between the two conditions; asterisks mark bands that are unique to one condition. Sizes of the molecular weight standards are indicated. Gels are representative of at least two independent experiments; similar results were obtained with subtilisin (see Fig. S2).

We next used a protease protection assay to test for any conformational changes in the purified proteins upon the addition of liposomes. In the presence of a low (1 μg/ml) concentration of trypsin, full-length PLIN3 showed fast degradation in solution, but was strongly stabilized by the addition of diphy liposomes (Fig. 2G). This is in agreement with previous work showing that the N-terminal half of PLIN3 is disordered in solution and folded upon binding to a lipid surface, thus decreasing its accessibility to proteolytic cleavage (8, 12). By contrast, the main PLIN3 degradation product, corresponding in size to PLIN3-4HB, showed stability over time (30 min) under both reaction conditions. A stable band of the same size was observed with purified PLIN3-4HB, confirming that this band corresponds to the 4HB region of PLIN3 (Fig. 2H). Interestingly, some weaker degradation bands also appeared below the PLIN3-4HB band, and the pattern of these bands was affected by the presence of liposomes. In particular, the number of degradation bands was decreased in the reaction with liposomes at longer time of protease treatment (30 min). This suggests that PLIN3-4HB is largely folded and protected from trypsin in the presence or absence of liposomes, but that it likely undergoes some conformational changes upon liposomes binding and becomes less accessible to proteolytic cleavage, which could be due to a decreased exposure of the αβ- subdomain. Similar results were obtained with another protease, subtilisin (Fig. S3).

In contrast to PLIN3-4HB, PLIN4-4HB did not show any lower-weight degradation bands when exposed to trypsin, and its susceptibility to trypsin was not affected by the addition of liposomes (Fig. 2I). This result is in agreement with the results of the sucrose flotation assay, confirming that PLIN4-4HB cannot bind to liposomes. Overall, PLIN4-4HB appeared less stable than PLIN3-4HB as it was largely degraded by 30 min of incubation with trypsin or subtilisin (Fig. 2I and S2C).

### PLIN3 4HB, but not PLIN4 4HB, can bind to LDs in cells

We next tested whether the 4HB domains from the two perilipins can bind to LDs in cells. First, we used as a model the budding yeast, which was shown previously to reproduce the interactions of human perilipins with LDs (41). We expressed PLIN3-4HB and PLIN4-4HB as GFP fusions in yeast cells lacking the yeast perilipin-like protein Pln1 (Pet10) (42) and grown to stationary phase in media supplemented with oleic acid (OA). These conditions were previously shown to promote maximum binding of perilipin AH regions, such as PLIN3-AH (see Fig. 1A), to LDs (13). For comparison, we also expressed full length PLIN3. Under these growth conditions, PLIN3-4HB efficiently localized to LDs, similar to PLIN3 and PLIN3 AH (Fig. 3A,B). By contrast, PLIN4-4HB was cytosolic in all cells (Fig. 3A,B).

**Figure 3.**
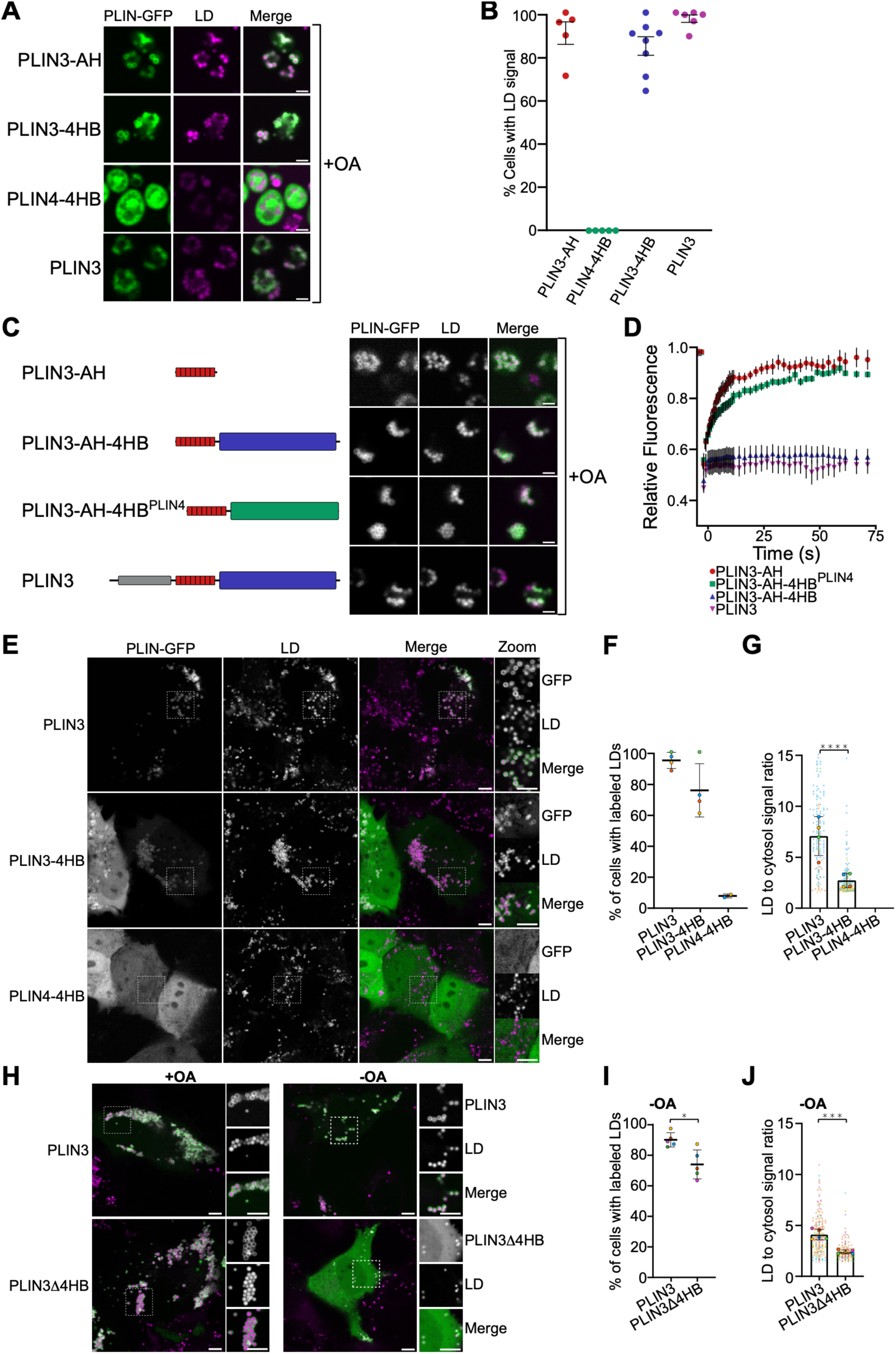
PLIN3 4HB, but not PLIN4 4HB, localizes to LDs in cells. **(A)** Localization of PLIN3-4HB, PLIN4-4HB and of the repetitive AH region of PLIN3 (PLIN3-AH), all fused to GFP, in budding yeast grown in the presence of oleic acid (OA) for 24h. LDs were stained with AUTODOT. Scale bar = 5 μm. **(B)** Quantification of the percent of cells with GFP signal on LDs from experiments shown in A). Each dot represents around 100 cells from a single experiment; mean ± SEM **(C)** Localization of PLIN3-AH, a truncated PLIN3 containing only the AH and 4HB regions (PLIN3-AH-4HB), a chimera protein where PLIN3-AH was fused to PLIN4-4HB (PLIN3-AH-4HB^PLIN4^), and full-length PLIN3, all fused to GFP, in budding yeast grown in the presence of oleic acid (OA) for 24h. LDs were stained with AUTODOT. Scale bar = 5 μm. **(D)** FRAP analysis of the dynamics of the constructs shown in (C) at the surface of LDs in yeast grown in the presence of OA. Curves from 10 to 20 independent measurements are shown, with error bars showing SD. **(E)** Localization of PLIN3, PLIN3-4HB and PLIN4-4HB upon transient expression in HeLa cells grown in media supplemented with OA. All proteins contain a C-terminal GFP tag. LDs were stained with Lipi-Blue. Magnifications (1.5x) of the indicated areas are shown on the right. Scale bars: 5 µm and 2µm in zoom-ins. **(F, G)** Quantification of experiments shown in panels E, showing % of cells with protein signal on LDs (F) and the ratio of GFP signal between LDs and the cytosol (G). Data points represent independent experiments (20-30 cells per experiment), each color representing a single experiment, lines show means or means ± SD from 2 to 5 experiments. **(H)** Localization of PLIN3 and PLIN3Δ4HB, lacking the 4HB domain upon transient expression in HeLa cells grown in media supplemented or not with OA. All proteins contain a C-terminal GFP tag. LDs were stained with Lipi- Blue. Magnifications (1.5x) of the indicated areas are shown on the right. Scale bars: 5 µm and 2µm in zoom-in’s. **(I, J)** Quantification of experiments shown in (H), showing % of cells with protein signal on LDs as in (F), or the ratio of GFP signal between LDs and the cytosol as in (G). Small symbols represent the mean of ratios for LDs in one cell (single z-section), large symbols represent the mean of each independent experiment, different colors are used for different experiments. Lines show means ± SEM from 5 independent experiments. Samples were compared using a Mann-Whitney test, *P <0.05, ***P<0.001, ****P<0.0001.

We then used fluorescence recovery after photobleaching (FRAP) to probe the dynamics of the interaction of various constructs with LDs (Fig. 3C,D). Whereas the PLIN3-AH region displayed a dynamic association with LDs, as previously shown (13), full length PLIN3 or a construct containing the two domains of PLIN3 (PLIN3-AH-4HB) showed very stable binding, reflected by a lack of recovery of the majority of the fluorescent signal on LDs (Fig. 3C,D). By contrast, the chimeric construct PLIN3-AH-4HB^PLIN4^, where PLIN3-AH was fused to PLIN4-4HB, displayed a dynamic association with LDs, similar to that of PLIN3 AH alone (Fig. 3C,D). This result suggests that even when PLIN4-4HB is brought close to the LD surface via AH targeting, it does not participate in LD binding.

We also evaluated the binding of PLIN3-4HB and PLIN4-4HB to LDs in HeLa cells. In agreement with the results with purified proteins and the budding yeast model, PLIN3-4HB partitioned between LDs and the cytosol, whereas full length PLIN3 was highly enriched on LDs. PLIN4- 4HB remained cytosolic in all cells (Fig. 3E-G). Furthermore, a truncated PLIN3Δ4HB protein, lacking the 4HB domain, displayed reduced binding to LDs compared to PLIN3 (Fig. 3H-J). These results confirm that the PLIN3-4HB domain contributes to LD binding, whereas PLIN4- 4HB does not.

### The αβ-subdomain in PLIN3 4HB contributes to its binding to LDs, but is not required for bundle stability

We wanted to understand the molecular basis for the difference in the behavior of the 4HB domains from PLIN3 and PLIN4. Close comparison of the two structural models shows that the main differences are at the base and at the top of the bundle structures. The αβ- subdomain in PLIN3-4HB is composed of disordered loops in PLIN4-4HB, and PLIN3 contains an additional predicted amphipathic α-helix, termed here helix-5 (H5), at the top of the bundle connecting helices 1 and 2; this α-helix is largely missing in the PLIN4-4HB structural model (Fig. 4A, B). Kymographs from MD simulations show that the features of the αβ-subdomain in PLIN3-4HB remain stable during a 1 μs simulation, as does the H5 (Fig. 4C, D). These features can also be observed in the kymograph of PLIN5-4HB, although the H5 is shorter. PLIN2-4HB may likewise contain an additional helix at the top of the bundle, whereas the N- and C-terminal segments surrounding the bundle are less structured. Kymograph of the MD simulation of PLIN4-4HB likewise shows that the regions corresponding to the αβ-subdomain in PLIN3-4HB are less structured and less stable during the 1 μs simulation (Fig. 4A).

**Figure 4.**
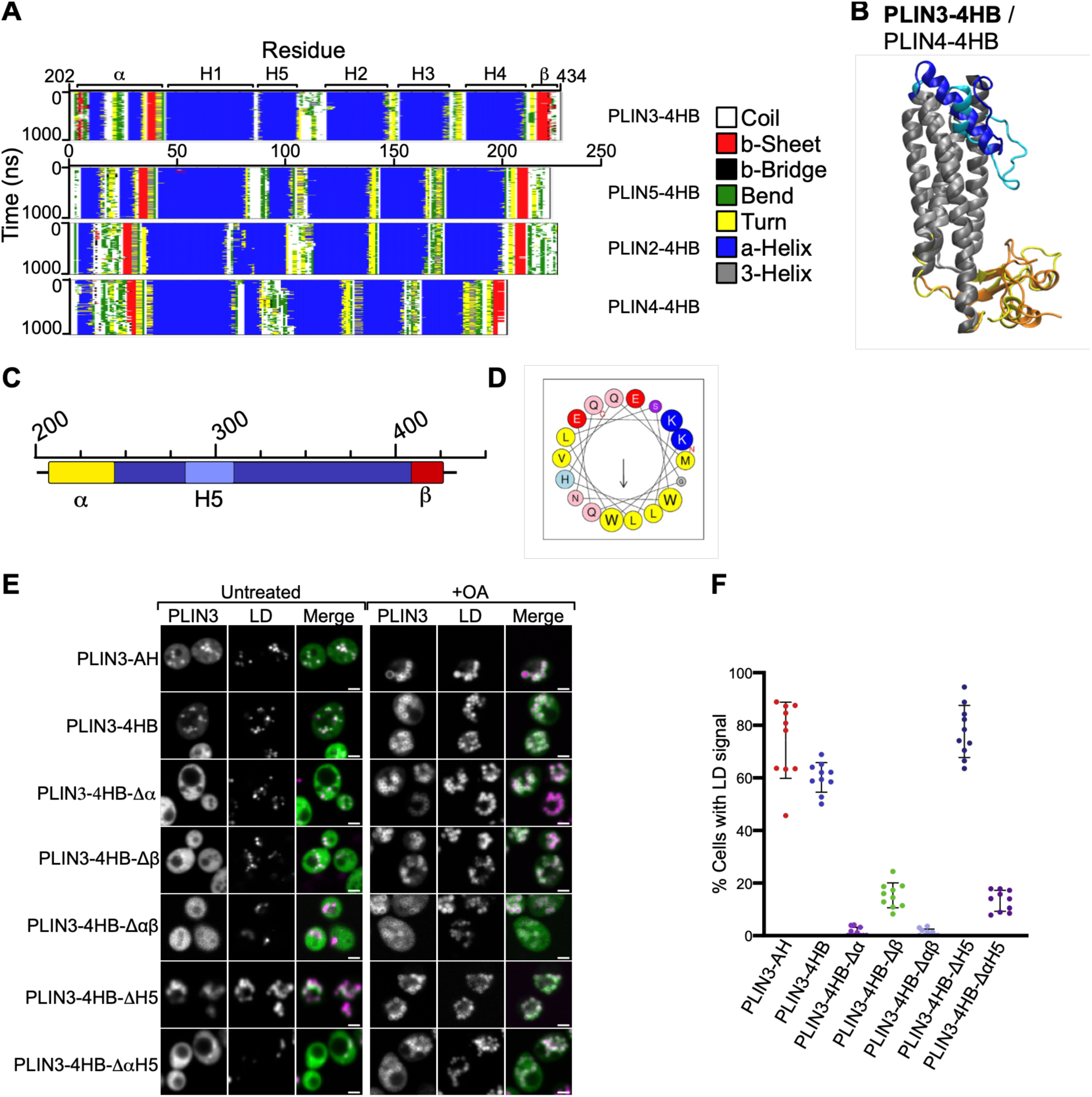
An αβ-lock is present in PLIN3 and PLIN5 4HB, but not in PLIN1, PLIN2 or PLIN4- 4HB region. (**A**) Kymographs from all-atom MD simulations (supp videos 1-5) of 4HB homology models from human PLIN3, PLIN5, PLIN2 and PLIN4, analyzed over 1 μs, showing the stability of different secondary structure elements along the amino acid sequence (x-axis) over time (y-axis). α- helices are shown in blue and β-sheets in red. Coil, β-bridge, bend, turn, and 3-helix are shown in white, black, green, yellow and grey, respectively. The four α-helices (H1, H2, H3 and H4), forming the predicted 4HB, and the additional α-helix (H5), are indicated, as well as the two parts of the αβ- subdomain (α and β). **(B)** Superposition of the homology models of PLIN3-4HB and PLIN4-4HB, highlighting the differences between the two structures: the αβ-subdomain and the additional helix H5 are present only in PLIN3-4HB. **(C)** Diagram of PLIN3 4HB sequence indicating the position of different deletions (α, β and H5). **(D)** Helical wheel representation of H5 from human PLIN3 4HB (aa 288-306). Arrow represents the hydrophobic moment. **(E)** The indicated PLIN3-4HB variants, tagged with GFP, were expressed in budding yeast and their localization was assessed by confocal microscopy under two growth conditions: 24 h in minimal media (’Untreated’), or 24h in media supplemented with OA (’OA’). LDs were stained with AUTODOT. Scale bar: 5 μm. **(F)** Quantification of results shown in (E), ’Untreated’ condition, showing the percentage of cells with LD-localized GFP signal. Each individual dot corresponds to a technical replicate with around 100 cells counted for each data point. Error bars represent ± 1SD.

To test whether the αβ-subdomain or the additional helix H5 in PLIN3-4HB contribute to its binding to LDs, we prepared mutants with deletions of these segments and analyzed their localization in budding yeast under two growth conditions: in glucose-containing media or after addition of oleic acid (OA) (Fig. 4E). Under the first growth condition, the deletion of the N-terminal (Δα), and to a slightly lesser extent the C-terminal (Δβ) portions of the αβ- subdomain strongly reduced the localization of PLIN3 4HB to LDs, whereas deletion of helix H5 did not significantly affect LD binding (Fig. 4E,F). In media supplemented with OA, all constructs could be observed on LDs, indicating that under the most permissive conditions for LD binding, the deletion mutants can still localize to LDs. However, the Δαβ mutant showed a high amount of cytosolic signal under these conditions (Fig. 4E, right columns).

We then purified the PLIN3-4HBΔα from *E. coli* in the same manner as we purified PLIN3-4HB, and tested its binding to liposomes. Even with the most permissive liposome composition (Diphy-50% PS), we could not detect any protein in the top (liposome-containing) fraction. The same result was obtained when PS concentration was reduced to 10% (Fig. 5A-B). These results complement the results obtained in yeast cells and suggest that the αβ-subdomain promotes PLIN3-4HB-binding to LDs.

**Figure 5.**
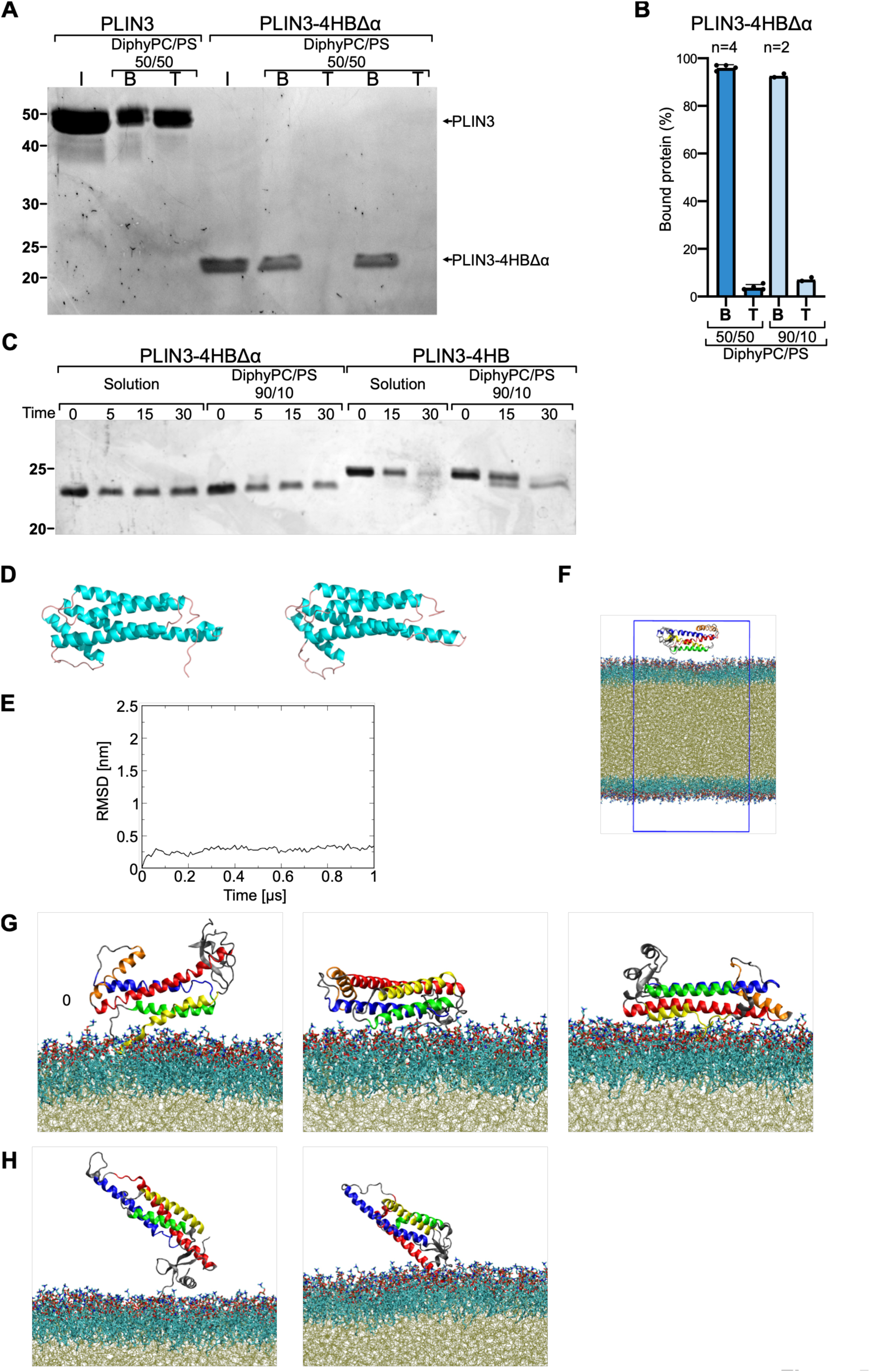
Mutagenesis of PLIN3 4HB shows that the αβ-subdomain contributes to LD binding, but is not required for 4HB stability. (A) PLIN3-4HBΔα mutant was purified from E. coli and its binding to Diphy liposomes (50% PC, 50% PS) was assessed by flotation on sucrose gradients. PLIN3 was used as control. Input (I) and fractions from the gradient (B, bottom; T, top) were analyzed by SDS-PAGE followed by SyproOrange staining. (B) Quantification of PLIN3-4HBΔα binding to DiPhy liposomes containing 50% or 10% PS. The number of replicates for each experiment is indicated above the bar graphs and liposome composition bellow the graph. (C) Limited proteolysis of Plin3-4HB and Plin3-4HBΔα (both at 0.1 mg/mL) in solution or in the presence of DiphyPC/PS 90/10 liposomes (1:100 protein-to-liposome ratio). The samples were incubated for the indicated time (0, 5, 15 or 30 min) with 0.2 µg/ml subtilisin and analyzed by SDS-PAGE using silver nitrate staining. Data is representative of two independent experiments. (D) Snapshots from the start (left image) and end (right image) of a 1 μs MD simulation of PLIN3-4HBΔαβ (Q233-W416, structure obtained by homology modeling) in solution, demonstrating the stability of this structure (see also Video S11). Similar results were obtained with the AlphaFold model (Video S12). (E) The root mean square deviation (RMSD) of the whole structures from the simulation shown in (D) is plotted over time; for comparison with wild-type PLIN4-4HB, see Fig. 1E. (F) Example of starting conformation for PLIN3-4HB in LD system. Phospholipids are colored in blue, triolein molecules are colored in tan. The hydrogens atoms and waters molecules are removed from the picture for clarity. (G) PLIN3-4HB was placed above the LD surface in three different starting positions and MD simulations were run for 500 ns. Images show the final positions from the three simulations. The four helices of the 4HB are colored in red (H1), blue (H2), green (H3) and yellow (H4). The extra-helix in PLIN3-4HB named H5 is colored in Orange. (H) PLIN4-4HB was placed above the LD surface in three different starting positions and MD simulations were run for 500 ns. Images show the final positions from two of the three simulations. In the third simulation (not shown), PLIN4-4HB became trapped between the two boundary conditions, creating an artefact. The four helices of the 4HB are colored as in (G). Movies of the simulations represented in (G; Video S12- S14) and (H; Video S15-S17) are shown in supplemental materials.

It was previously suggested that the 4HB of PLIN3 participates in the interaction with LDs owing to 4HB opening up on the LD surface, leading to the individual helices of the bundle laterally interacting with the LD surface (31). Furthermore, it was suggested that the αβ- subdomain may function to stabilize the 4HB structure. In support of this model, a small increase in LD binding can be observed with C-terminally truncated full-length PLIN3 (PLIN3Δβ) (31) (Fig. S4). These results contrast with the decrease in LD binding that we observe with PLIN3-4HB mutants, but can also be explained by synergistic effects between different parts of PLIN3 in LD/lipid surface binding (24). To further analyze the behavior of the 4HB domain, we evaluated the stability of folding of the purified αβ-subdomain mutant PLIN3- 4HBΔα by limited proteolysis in the presence or absence of diphy liposomes. Importantly, PLIN3-4HBΔα was even more resistant to proteolysis than wild-type PLIN3-4HB (Fig. 5C). Furthermore, the protease sensitivity of the PLIN3-4HBΔα mutant was not affected by the presence of Diphy liposomes, in accordance with its lack of binding to diphy liposomes or to LDs. The stability of the truncated PLIN3-4HBΔαβ mutant, lacking the full αβ-subdomain, can also be observed in MD simulations in solution, where it remained stable during the course of a 1 μs simulation for two models (AlphaFold or homology modeling) (Video S10 and S11); this can be observed in Fig. 5D, showing the starting and the final snapshot from the simulation, in which the structure of PLIN3-4HBΔαβ remained largely unchanged. The RMSD plot (Fig. 5E) shows very small fluctuation during the simulation (compared to RMSD obtained for PLIN 4HB domains in Fig. 1E).

To further explore the difference in the PLIN3-4HB and PLIN4-4HB LD affinities, we performed all-atom MD simulations in the presence of an LD surface. At the beginning of each simulation, we placed the protein above the LD, composed of triolein and palmitoyl-oleoyl-PC (POPC). The density of the POPC monolayer was adjusted such that the lipid packing was reduced to 80% of density in a POPC bilayer, introducing shallow lipid packing defects and an increase in surface tension, which promotes binding of purified PLIN3 to artificial LDs (22, 43, 44) (Fig. 5F). Simulations of 500 ns were repeated three times for both PLIN3-4HB and PLIN4-4HB starting with different orientations of the proteins compared to the LD surface. During the course of the simulations, PLIN3-4HB quickly (within the first 50-100 ns) approached the LD surface and remained at a close distance from the surface (Video S12-S14). By contrast, PLIN4- 4HB oscillated above the surface and did not stabilize with respect to its distance from the LDs, except for an interaction via carboxy-terminal residue, which may be due to unprotected positive charge (Video S15-S16). In the final simulation, PLIN4-4HB became trapped between two boundary conditions, therefore this simulation was excluded from further analysis (Video S17). The final positions of PLIN3-4HB and PLIN4-4HB in the simulations are shown in Fig. 5G and H. The three simulations for PLIN3-4HB did not converge to a similar LD-bound orientation and did not reveal that the αβ-subdomain directly interacts with the LD surface. Longer MD simulations or possibly an alternative force-field (45) might be needed to better capture the most relevant PLIN3-LD interaction and should be extended in future work.

## Discussion

Perilipins are the most abundant LD surface proteins in most cell types and have long been recognized to play an important role in cellular LD homeostasis. They directly interact with the lipid surface of LDs; however, their structural organization on the LD surface is still poorly understood. In this work, we have addressed the behavior of the C-terminal 4HB domain. We show that this folded domain is present in all human perilipins except PLIN1. Whereas the solution structure of the 4HB from PLIN3 was first determined in 2004, the function of this domain on the LD surface has remained mysterious. Here, we show that the 4HB of PLIN3, but not of PLIN4, can directly interact with the LD surface. The PLIN4-4HB lacks a stably-structured αβ-subdomain that is present in PLIN3-4HB, and the αβ-subdomain is required for the interaction of PLIN3-4HB with the LD surface. Furthermore, we show that the αβ-subdomain is not required to maintain the stability of the helical bundle, as determined by limited proteolysis and supported by MD simulations. We therefore suggest that the αβ-subdomain region may directly interact with the LD surface, thus further stabilizing PLIN3 interaction with LDs. Modeling and MD simulations suggest that a stable αβ-subdomain may also be present in PLIN5. In PLIN4, which arose in mammals by gene duplication from *PLIN5,* the LD-interacting repetitive AH region has greatly expanded (5, 13), and the 4HB domain may have concomitantly lost the ability to promote LD binding.

Another important conclusion from this work is that the 4HB of PLIN3, as well as those of PLIN4 and PLIN5, are over-all stable structures that do not show a tendency to undergo large conformational changes, such as bundle opening, to permit the interaction of these helices with the LD surface. This mechanism was previously suggested for the 4HB of PLIN3 based on its structural similarity with apolipoproteins, which undergo large conformational changes upon interaction with lipoprotein particles (31, 35). We show instead that the 4HB of PLIN3 is a stable structure and that the αβ-subdomain is not required to maintain its stability. The 4HB of PLIN3 interaction with the lipid surface of LDs could promote fatty acid channeling, as has been suggested for PLIN2 and PLIN5 4HBs, which can accommodate specific fatty acids within their hydrophobic cleft (29, 37). A recently-published study suggests that the hydrophobic cleft of the 4HBs from PLIN2, PLIN3, PLIN4 and PLIN5 can accommodate phosphatidylethanolamine (PE), which would target these proteins towards PE-rich LDs (46). However, no PE-specificity has been observed with full-length perilipins, and it is also difficult to reconcile the binding of this phospholipid deep within the 4HB structure with its proposed role in LD surface targeting. Finally, the 4HB domain of perilipins could also participate in protein-protein interactions, including protein oligomerization on the LD surface; this could explain why the binding of the PLIN3-4HB-containing constructs to LDs in budding yeast is so stable, in contrast to the dynamic binding of the PLIN3-AH. These questions remain to be explored in future work.

## Experimental procedures

### Protein modeling

We generated models of the C-terminal (4HB) homology regions from the human perilipins PLIN1-PLIN5 using AlphaFold3 website (AF) (47) using the human UniprotDB sequences. We selected the best structure (from 5 generated) for each model.

For Homology modeling, we used Modeller tool (48) to generate homology models based on the structure of mouse PLIN3-4HB (33). The region 289-320 was not been resolved in the crystal structure of the mouse PLIN3-4HB and this part has been considered as a gap in the alignment sequences. For each 4HB PLINs, we generated 100 models from Modeller and selected the best energy function model. The images have been generated with the visualization software PyMOL (Schrödinger LLC. (n.d.). PyMOL Molecular Graphics System (2.5)).

### Molecular dynamics simulations

The structural models (AF and Modeller) were equilibrated during 1 µs all-atom MD simulations in solution using the Charmm36 force-field (49) and GROMACS tool version 2023.4 (50). After minimization process, we added water molecules and neutralized the system with 120 mM Na^+^ and Cl^-^ ions. A cut-off distance of 1.2 nm was used for generating the neighbor list and this list was updated at every step. Long-range electrostatic interactions were calculated using the particle mesh Ewald summation methods (PME). Periodic boundary conditions were used. During the production run, the V-rescale thermostat and C-rescale barostat stabilized the temperature at 310 K and pressure at 1 bar, respectively.

To perform simulations of protein binding to an LD surface, we constructed LD models as described in (Araújo et al., 2024). The triolein (TG(18:1/18:1/18:1)) topology was modified as in (52). We started from bilayers (14.1 × 14.1 nm) containing 412 molecules of PC(16:0/18:1) in water (50 Å ) obtained and equilibrated with Chamm-Gui tools (53). We incorporated 864 molecules of triolein between the two monolayers and then performed minimization and equilibration for 110 ns using GROMACS 2023.4. After equilibration, we removed 20% of phospholipids to obtain 80% PC coverage (330 phospholipids). A cut-off distance of 1.2 nm was used for generating the neighbor list and this list was updated at every step. Long-range electrostatic interactions were calculated using the particle mesh Ewald summation methods (PME). Periodic boundary conditions were used. During the production run, the V-rescale thermostat and C-rescale barostat stabilized the temperature at 310 K and pressure at 1 bar, respectively. To maintain the LD-water systems, we used a compressibility of 0 on x/y axis in semi-isotropic condition. The simulations were performed for 500ns for each system and coordinates were saved every 100 ps. Three independent replicas were performed for each system by changing the orientation of the PLIN3-4HB or PLIN4-4HB in water above the LD surface at the start of each simulation.

The MD analyses of density profiles were performed using GROMACS utilities. The movies has been generated with VMD software (54).

### Plasmid DNA construction

All plasmids used in this study are listed in Supplementary table S1. The plasmid pGREG576 for expression of full-length human PLIN3 from an ADH1 promotor (gift from R. Schneiter, U. of Fribourg) (41) was used as the backbone to construct all yeast expression plasmids. A point mutation in the PLIN3-encoding sequence in this plasmid was corrected by site-directed mutagenesis (R264G) to obtain the PLIN3 sequence matching Uniprot: O60664. truncation mutants were generated by linearisation of the corrected pGREG576 by PCR, followed by circularisation of the plasmid using T4 DNA ligase (NEB). The PLIN4-4HB-encoding sequence was amplified from a synthetic gene encoding E. coli codon-optimized PLIN4- sequence from Eurofins Genomics (https://www.eurofinsgenomics.eu) and cloned into linearized pGREG576 backbone via NEBuilder HiFi-assembly (New Egland Biolabs), either in fusion with GFP or in fusion with the AH region of PLIN3 and GFP.

To construct plasmids for protein expression in human cells, PLIN3-encoding sequences were amplified from the mEGFP-PLIN3 Addgene plasmid # 222426 (gift from Stephen Roy, Downie et al., 2025), or a PLIN4-encoding human codon-optimized synthetic gene from Eurofins Genomics. They were inserted into the pEGFP-C1 plasmid (Clontech), linearised with EcorRI using NEBuilder HiFi-assembly (New Egland Biolabs).

For expression of proteins in *E. coli*, PLIN3-4HB sequences were amplified from the pET16b- His-PLIN3 plasmid (22), and the PLIN4-4HB sequence was amplified from human cDNA (gift from A. Ruggieri, Fondazione IRCCS Istituto Neurologico Carlo Besta, Milano). They were then cloned into the pET16b-GST-Tev vector, which contains an N-terminal GST tag followed by a TEV cleavage site, using *NcoI* and *BamHI* restriction sites.

All PCR amplifications were carried out using high-fidelity enzyme Herculase DNA Polymerase (Ozyme). All plasmid sequences were verified by Sanger sequencing.

### Protein purification

All proteins were purified from *E. coli*. Plin3-4BH ,PLIN3-4HBΔα and Plin4-4HB were purified using a protocol adapted from the protocol used for the purification of mouse PLIN3-4HB (33). ArticExpress *E. coli* competent cells (Agilent) transformed with expression plasmids were grown to O.D. ≈ 0.6 at 37°C from a liquid preculture and induced with 0.1 mM IPTG overnight at 16 °C. Cells from 0.5 l cultures were collected by centrifugation and frozen. The bacterial pellets were thawed in lysis buffer (50 mM Tris-HCl pH 7.5, 150 mM NaCl, 1 mM DTT), supplemented with 0.5 mM PMSF, 0.1 mM pepstatin, 0.02u/L of benzonase endonuclease and Complete protease inhibitor cocktail (Roche). Cells were broken by sonication. The lysate was centrifuged at 100,000 × g for 30 min at 4°C. Supernatant was loaded on 1ml Glutathione Sepharose beads (Cytiva), equilibrated with purification buffer B1 (20 mM Tris-HCl pH 7.5, 10mM NaCl, 1mM DTT) and incubated with the beads for 2h at 4°C with gentle rotation. Beads were collected by centrifugation at low speed, and washed with buffer B1, then with buffer B2 (20 mM Tris-HCl pH 7.5, 10mM NaCl, 1mM TCEP). Beads were then incubated overnight at 4°C under gentle rotation in 5ml of buffer B2 containing TEV protease (95 µg/ml). The resulting supernatant was incubated at 4°C for 1h with pre-equilibrated Ni-NTA beads (QIAGEN) to remove TEV (containing a C-terminal His-tag); efficient removal of TEV was verified by SDS-PAGE. Pooled unbound fractions were concentrated on an Amicon 10kDa MWCO cell and further purified by size exclusion chromatography on a Superose 6 10/300 column or Hiprep 16/60 Sephacryl S-300 HR (Cytiva) equilibrated with buffer D (50 mM Tris-HCl pH 7.5, 120mM NaCl, 1mM TCEP). Fractions representing the monomeric protein peak were pooled, aliquoted and snap frozen for storage at -80°C. Protein purity was analysed by SDS-PAGE and the monomeric state and folded conformation of purified proteins were verified by dynamic light scattering (DLS) and by circular dichroism (CD) spectroscopy.

Full length human PLIN3 was purified as described in (22) and PLIN4-12mer (396-amino acid fragment of the repetitive region) was purified as described in (11).

### Liposome preparation

The following phospholipids used in the experiments, 1,2-dioleoyl-sn-glycero-3- phosphocholine (DOPC), 1,2-dioleoyl-sn-glycero-3-phospho-L-serine (DOPS), 1,2-diphytanoyl-sn-glycero-3-phosphocholine (diphy-PC), 1,2-diphytanoyl-sn-glycero-3-phospho-L-serine (diphy-PS) and Rhodamine-PE (L-α-Phosphatidylethanolamine-N-lissamine rhodamine-B - sulfonyl) were purchased from Avanti Polar Lipids as chloroform solutions. Lipids were stored as chloroform stock solutions at -20°C under argon.

Phospholipids in chloroform were mixed at the desired molar ratio and dried under argon flux. To prepare fluorescently-labeled liposomes, 0.1 % of Rhodamine-PE was included in the mixture. The lipid film was hydrated in 1 mL of HK buffer (50 mM HEPES, pH 7.4, and 120 mM K-Acetate) to a concentration of 2 mM phospholipid and freeze-thawed five times in liquid nitrogen. Prior to use, 500 µL of liposomes were extruded 19x through a polycarbonate filter of 0.1 µm pore size using a mini-extruder. Extruded liposomes were stored at room temp and used within 2 days following extrusion.

### Liposome flotation assay

Proteins (5 μM) were mixed with liposomes (500 μM) in HK buffer (50 mM HEPES, pH 7.4, and 120 mM K-Acetate) containing 1 mM MgCl_2_ and 1 mM DTT 5 µM in 120 µL final volume. The mix was incubated for 10 min at 25°C. 100 µL of each sample was then mixed with 140 µL of 60% sucrose (w/v) in HKM buffer to on obtain a final sucrose concentration of 35%, transferred into thick-wall centrifugation tubes (Beckman), and overlayed with 200 μL of 25% sucrose in HKM buffer, followed by 50 μL of HKM buffer. Gradients were centrifuges at 240,000 × g, 20°C, for 1 hour in an S55-S spin-out rotor. Following centrifugation, three fractions were collected from the bottom and labelled bottom (250 μL), middle (140 μL), and top (100 µL) and analyzed by SDS-PAGE. Total fractions from each mix were included for quantification. Protein staining was performed using SyproOrange staining (ThermoFisher), and protein bands were quantified using ImageJ.

### Protease protection assay

Prior to the experiment, proteins were centrifuged 10 min at 20,000 g, 4°C, to remove any precipitate, and diluted in HK buffer (50 mM HEPES, pH 7.4, and 120 mM K-Acetate) containing 1 mM MgCl_2_ to 0.1 mg/mL. was mixed or not with diphytanoyl (50% PC, 50% PS) liposomes at 1:100 protein-to-phospholipid molar ratio in 120 µL final volume. The mix was incubated for 10 min at 25°C with 300 rpm agitation prior to addition of 0.2 µg/mL of subtilisin or 1 µg/mL trypsin. At indicated times, 20 μL of the reaction was removed and the reaction was stopped immediately with 2mM PMSF. Samples were analyzed by SDS-PAGE and proteins were stained with SyproOrange or with Silver stain.

### Circular dichroism

CD measurements were conducted on a Jasco J-815 spectrometer at room temperature with a quartz cell of 0.05 cm path length. Each spectrum is the average of several scans recorded from 200 to 260 nm with a bandwidth of 1 nm, a step size of 0.5 nm and a scan speed of 50 nm min^−1^. The buffer used was Tris 10 mM, pH 7.5, KCl 150 mM. Ellipticity was converted to mean residue ellipticity (MRE) by dividing by the product of protein concentration, residue number and path length distance.

### Yeast growth and media

Yeast strains used were: BY4742 MATα *his3Δ1 leu2Δ0 lys2Δ0 ura3Δ0 (Euroscarf)*, and BY4742 *pln1Δ::KANMX4 (Euroscarf)* (56). Yeasts were transformed by standard lithium acetate/polyethylene glycol procedure with low-copy plasmids expressing different PLIN3/PLIN4-GFP constructs under the control of ADH1-promotor. Yeast cells expressing different constructs were grown in synthetic complete medium lacking uracil (SC-Ura, 6.7 g/l yeast nitrogen base, amino acid supplement without uracil, 2% glucose). To induce LDs, yeast cells were either grown in SC-Ura for 24h at 30°C (stationary phase) or for 24h in SC-Ura, followed by 24h incubation in oleic acid (OA) medium (0.67% yeast nitrogen base without amino acids, 0.1 % yeast extract, 0.1 % (v/v) oleate, 0.25 % (v/v) Tween 40, amino acid supplement lacking uracil).

### Cell culture and transfection

HeLa cells were grown in Dulbecco’s modified Eagle’s medium (DMEM) supplemented with 4.5 g/l glucose (Thermo Fisher), 10% fetal bovine serum (FBS, Thermo Fisher), 0.584g/L L-Glutamine (Thermo Fisher) and 1% Penicillin/Streptomycin antibiotics (Thermo Fisher). For protein expression, subconfluent cells were transfected with FuGENE HD transfection reagent (Promega) in Optimem medium (Thermo Fisher) for 5 h, followed by 18 h of growth in standard growth medium without phenol red. When indicated, the cell medium was supplemented with 200 µM OA (O1383; Sigma-Aldrich) complexed with fatty acid-free BSA (Sigma-Aldrich).

### Fluorescent microscopy

Yeast cells were harvested by centrifugation, washed, labelled with Autodot blue dye (Clinisciences) diluted 1000-fold for 30 min at room temp, after which the cells were washed twice and imaged.

Transfected HeLa cells were stained with LipiBlue (Dojindo) diluted 1000-fold and Cellmask Deep red (Thermo Fisher) diluted 1000-fold at 37°C. Cell were washed with PBS and imaged at 37°C in glass bottom plates (GREINER BIO ONE, 627870) in standard growth medium without Phenol red with an inverted Olympus IXplore Spin SR microscope coupled with a spinning disk CSU-W1 head (Yokogawa) using 60X UPLXAPO 1.42 NA DT 0.15mm oil- immersion objective piloted with the Cellsens software. Images were processed with ImageJ and prepared for figures with Affinity (Canvas X).

### Fluorescence recovery after photobleaching (FRAP)

FRAP assays in yeast cells were performed on yeast grown to exponential growth phase and then treated for 24h with Oleic acid on glass slides using the OLYMPUS SR spinning disk microscope and 60x objective, bleaching laser with a wavelength of 488 nm and CellSens software. A circular area comprising a single isolated LD (about 15pixels of diameter) in a cell expressing a GFP-fusion protein was bleached. 5 images were taken before bleaching, followed by a post-bleach time-course: 12 s of 1 image/0.6s, 50 s of 1 image/2.5 s, 100 s of 1 image/5 s, and 200 s of 1 image/10 s. Background fluorescence outside the cell was subtracted from the bleached area and the signal was normalized to the whole cell signal for each time- point. Data was processed with Excel and plotted using GraphPad Prism 10 (GraphPad Software, Boston, Massachusetts USA).

### Image analysis

Images were analysed using ImageJ/Fiji (57). To quantify the number of yeast cells with protein signal on LDs, cells were counted manually after applying the same brightness/contrast settings to all images.

To determine PLIN4 binding to LDs in HeLa cells, a single z-section that contained the most LDs in a cell was analysed using a custom image-processing pipeline. Plasma membrane channel (CellMask deep red) was used to segment individual cells with Cellpose 4.0.6 (58). After segmentation, images were processed through a custom FIJI macro were; cells with a mean signal in the GFP channel between 1.4 and 10 times the background were quantified. LDs in the selected z-section were identified using the LipiBlue dye signal and segmented with a custom Labkit classifier (59). The ’Connected components labelling’ tool (MorphoLibJ) (60) was used to identify individual LDs. To determine the ratio between LD-bound and soluble GFP fluorescent signal, intensity values were extracted along line profiles drawn across each LD (10 regularly-spaced lines per LD). The LD-bound GFP fluorescence intensity was measured for 5 pixels on either side of the LipiBlue signal along each line, and the cytoplasmic fluorescence intensity was determined up to 15 pixels away from the LipiBlue signal; signals overlapping with LipiBlue labelling from neighbouring LDs were excluded. The mean bound and cytoplasmic signals were recorded for each line, and the mean values for all 10 lines were used to calculate the bound-to-cytoplasmic signal ratio for each LD.

LDs with a GFP signal ratio > 1.4 were considered positive, and the percentage of positive LDs per cell was determined as the number of positive LDs relative to the total number of identified LDs. To determine the percentage of cells with LD signal, we noted that most cells displaying protein binding to LDs presented between 85% and 100% of labelled LDs, whereas cells with low signal showed a lower percentage of labelled LDs. To include such cells in the quantification, a cell was considered as presenting LD-bound protein if at least 20% of its LDs were positive. For graphical representation of the LD-to-cytosol signal ratio, only positive LDs were considered, and a mean ratio value was determined for each cell. Data were processed with Excel and plotted using GraphPad Prism 10 (GraphPad Software, Boston, Massachusetts, USA).

### Statistical analysis

All statistical analyses were performed using GraphPad Prism (ver. 10.6.1) software (GraphPad Software, Boston, Massachusetts, USA). For cross-analysis, a Shapiro test was used to control for normality of the samples, followed by a pair-wise t-test or a Mann-Whitney test. Details are described in the respective figure legends.

## Data availability

All data are contained within the article and the supporting information.

## Supporting Information

This article contains supplemental information: Supplemental Table S1, Supplemental Figures S1 - S4 and Supplemental videos S1-S17.

## Acknowledgments

We thank C Franckhauser (CRBM, Montpellier) for technical support, and C Syska (CRBM, Montpellier), B Antonny (IPMC, Sophia Antipolis) for helpful discussions and critical reading of the manuscript. We acknowledge the imaging facility MRI, member of the national infrastructure France-BioImaging (https://ror.org/01y7vt929) supported by the French National Research Agency (ANR-24-INBS-0005 FBI BIOGEN), the joint IGMM-CRBM “yeast media and technologies service”, and the Synbio3 platform (IBMM, Montpellier University, France), supported by GIS IBISA. This work was supported by the European Research Council (ERC Synergy 856404, SPHERES), and by the Agence Nationale de la Recherche (ANR-23-CE44- 0026).

## Author contributions

Conceptualization (R.G., A.C.); data curation (C.M., R.G.); formal analysis (C.M., B.S., R.G.); funding acquisition (A.C.); investigation (C.M., B.S., C.F., N.F., R.G.); methodology (C.M., A.B., R.G., A.C.); project administration (A.C.); supervision (A.B., R.G., A.C.); visualization (C.M., B.S., R.G.); writing – original draft (A.C.); writing – review & editing (C.M., B.S., R.G., A.C.)

## Conflict of interest

The authors declare that they have no conflicts of interest with the contents of this article.

## Notes

### Competing Interest Statement

The authors have declared no competing interest.

