## Supplemental data for "Comparison of the 4-helix bundle domains of perilipin 3 and perilipin 4 identifies features that contribute to lipid droplet binding"

### Supplemental Information

Cyril Moulin[1‡], Bayane Sabbagh[1‡], Amel Bahloul[1,2], Nicolas Fuggetta[1], Romain Gautier[3*], Alenka Čopič[1*]

[1]Centre de Recherche en Biologie cellulaire de Montpellier-CRBM, Université de Montpellier, CNRS, UMR 5237, Montpellier, France.
[2]Institut des Neurosciences de Montpellier-INM, Université de Montpellier, INSERM U1298, Montpellier, France.
[3]Université Côte d'Azur, CNRS and Inserm, Institut de Pharmacologie Moléculaire et Cellulaire, UMR 7275, Sophia Antipolis France.

‡ These authors contributed equally to this work.

This file contains:

Supplemental Table S1, Supplemental Figures S1 - S4 and Captions for Supplemental Videos S1-S17.

**Supplementary Table 1. Plasmids used in this study.**

| Name | Insert | Region (aa) (1) | Vector | Host (2) | Source |
|---|---|---|---|---|---|
| pGFP-Plin3 | Human (h) PLIN3 | Full cDNA | pGREG576 (ADH1pr, GFP) | Yeast | (41) |
| pCM90 | hPLIN3 | Correction G264 to R (3) | pGREG576 (ADH1pr, GFP) | Yeast | This study |
| pCM105 | hPLIN3-AH | PLIN3 (114-205) | pGREG576 (ADH1pr, GFP) | Yeast | This study |
| pCM107 | hPLIN3-4HB | PLIN3 (206-434) | pGREG576 (ADH1pr, GFP) | Yeast | This study |
| pCM123 | hPLIN3-4HBΔα | PLIN3 (244-434) | pGREG576 (ADH1pr, GFP) | Yeast | This study |
| pCM114 | hPLIN3-AH - hPLIN4-4HB | PLIN3 (114-205)-PLIN4 (1061-1371) | pGREG576 (ADH1pr, GFP) | Yeast | This study |
| pCM118 | hPLIN4-4HB | PLIN4 (1061-1371) | pGREG576 (ADH1pr, GFP) | Yeast | This study |
| pCM130 | hPLIN3-4HBΔH5 | PLIN3 (206-287,316-434) | pGREG576 (ADH1pr, GFP) | Yeast | This study |
| pCM131 | hPLIN3-4HBΔαΔH5 | PLIN3 (244-287,316-434) | pGREG576 (ADH1pr, GFP) | Yeast | This study |
| pCM134 | hPLIN3-4HBΔβ | PLIN3 (206-413) | pGREG576 (ADH1pr, GFP) | Yeast | This study |
| pCM135 | hPLIN3-4HBΔαβ | PLIN3 (244-413) | pGREG576 (ADH1pr, GFP) | Yeast | This study |
| pmEGFP-PLIN3-addgene222426 | hPLIN3 | PLIN3 (1-434) | pmeGFP-C1 | Mamm | (55) |
| pCM187 | hPLIN3 | PLIN3 (1-434) | pGFP-C1 (CMVpr, GFP) | Mamm | This study |
| pCM188 | hPLIN3Δ4HB | PLIN3 (1-205) | pGFP-C1 (CMVpr, GFP) | Mamm | This study |
| pCM191 | hPLIN3-4HB | PLIN3 (206-434) | pGFP-C1 (CMVpr, GFP) | Mamm | This study |
| pCM194 | hPLIN4-4HB | PLIN4 (1061-1371) | pGFP-C1 (CMVpr, GFP) | Mamm | This study |
| pET16b-His-PLIN3 | hPLIN3 | *E. coli* codon optimized PLIN3 (1-434) | pET16b-His10-Tev-LIC | *E. coli* | (22) |
| pET16b-GST-Tev | GOLPH3 | -- | pET16b-GST-Tev | *E. coli* | |
| pBS03 | hPLIN4-4HB | PLIN4 (1150-1371) | pET16b-GST-Tev | *E. coli* | This study |
| pBS08 | hPLIN3-4HB | PLIN3 (197-434) | pET16b-GST-Tev | *E. coli* | This study |
| pBS12 | hPLIN3-4HBΔα | PLIN3 (244-434) | pET16b-GST-Tev | *E. coli* | This study |
| pKE23 | hPLIN4-12mer | PLIN4 (524-919), 12 x 33-aa repeats | pET21b | *E. coli* | (11) |
| pCM189 | hPLIN3Δβ | PLIN3 (1-413) | pGFP-C1 (CMVpr, GFP) | Mamm | This study |
| pCM91 | hPLIN3 AH-4HB | PLIN3 (114-434) | pGREG576 (ADH1pr, GFP) | Yeast | This study |
| pCM92 | hPLIN3 PAT-4HB | PLIN3 (1-113/206-434) | pGREG576 (ADH1pr, GFP) | Yeast | This study |
| pCM93 | hPLIN3 PAT-AH | PLIN3 (1-205) | pGREG576 (ADH1pr, GFP) | Yeast | This study |

(1) Position of amino acids (aa) in human perilipin sequences.
(2) Mamm: mammalian cells
(3) All yeast PLIN3 expression vectors are derived from pGFP-Plin3 (41), in which a mutation at position 264 was corrected, G264->R, in accordance with the human PLIN3 Uniprot sequence.

### Supplement figures

### A

```
CLUSTAL multiple sequence alignment by MUSCLE (3.8)


PLIN2h          ---------LPLTEEELEKEAKKVEGFDLVQ------KPSYYVRLGSLSTKLHSRAYQQA
PLIN4h          --------FHPMNAEEQAQLAASQPGPKVLSAE----QGSYFVRLGDLGPSFRQRAFEHA
PLIN3h          ---------LPLTDAELARIATSLDGFDVASVQQQRQEQSYFVRLGSLSERLRQHAYEHS
PLIN5h          KSEELVDHFLPMTEEELAALAAEAEGPEVGSVEDQRRQQGYFVRLGSLSARIRHLAYEHS
                          *:.  *    * .  * .: .      : .*:****.*.  :.  *::::

PLIN2h          LSRVKEAKQKSQQTISQLHSTVHLIEFARKNVYSANQKIQDAQDKLYLSWVEWKRSIGYD
PLIN4h          VSHLQHGQFQARDTLAQLQDCFRLIEKAQQ--------APEGQPRLDQGSGASA------
PLIN3h          LGKLRATKQRAQEALLQLSQVLSLMETVKQGV---DQKLVEGQEKLHQMWLSWNQKQLQG
PLIN5h          VGKLRQSKHRAQDTLAQLQETLELIDHMQCGV---TPTAPACPGKVHELWGEWGQRP---
                :..:.  : .:.::: ** . . *::  .               .:

PLIN2h          DTDESHCAEHIESRTLAIARNLTQQLQTTCHTLLSNIQGVPQNIQDQAKHMGVMAGDIYS
PLIN4h          --EDAAVQEERDAGVLSRVCGLLRQLHTAYSGLVSSLQGLPAELQQPVGRARHSLCELYG
PLIN3h          PEKEPPKPEQVESRALTMFRDIAQQLQATCTSLGSSIQGLPTNVKDQVQQARRQVEDLQA
PLIN5h          --PESRRRSQAELETLVLSRSLTQELQGTVEALESSVRGLPAGAQEKVAEVRRSVDALQT
                   :.   .  :  .*    .: .:*: :   * *.:.*:*   :: .         :

PLIN2h          VFRNAASFKEVSDSLLTSSKGQLQKMKESLDDVMDYLVNNTPLNWLVGPFYPQLTESQNA
PLIN4h          IVASAGSVEELPAERLVQSREGVHQAWQGLEQLLEGLQHNPPLSWLVGPFA------LPA
PLIN3h          TFSSIHSFQDLSSSILAQSRERVASAREALDHMVEYVAQNTPVTWLVGPFAPGITEKAPE
PLIN5h          AFADARCFRDVPAAALAEGRGRVAHAHACVDELLELVVQAVPLPWLVGPFAPILVERPEP
                 . .  .. ::.   *....  :   .  :: ::: : :  *: ******

PLIN2h          QDQGAEMDKSSQETQRSEHKTH
PLIN4h          GGQ-------------------
PLIN3h          EKK-------------------
PLIN5h          LPD-------------------
```

### B

**Superposition of PLIN1 and PLIN3 C-terminal structures**

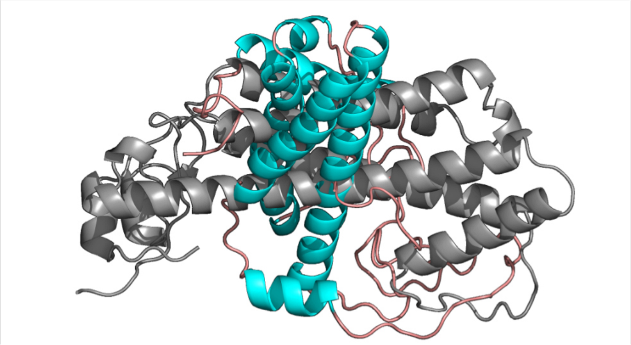

**FigureS1. Additional data related to Figure 1.**
**(A**) Alignment of sequences of the C-terminal regions of human PLIN2 (Q99541: L191-H437*), PLIN3 (O60664: L204-K434*), PLIN4 (Q96Q06: F1162-Q1371*, and PLIN5 (Q00G26: K157-D391. **(B)** Superposition of Alphafold 3 models of 4HB-homology regions from human PLIN1 and PLIN3.

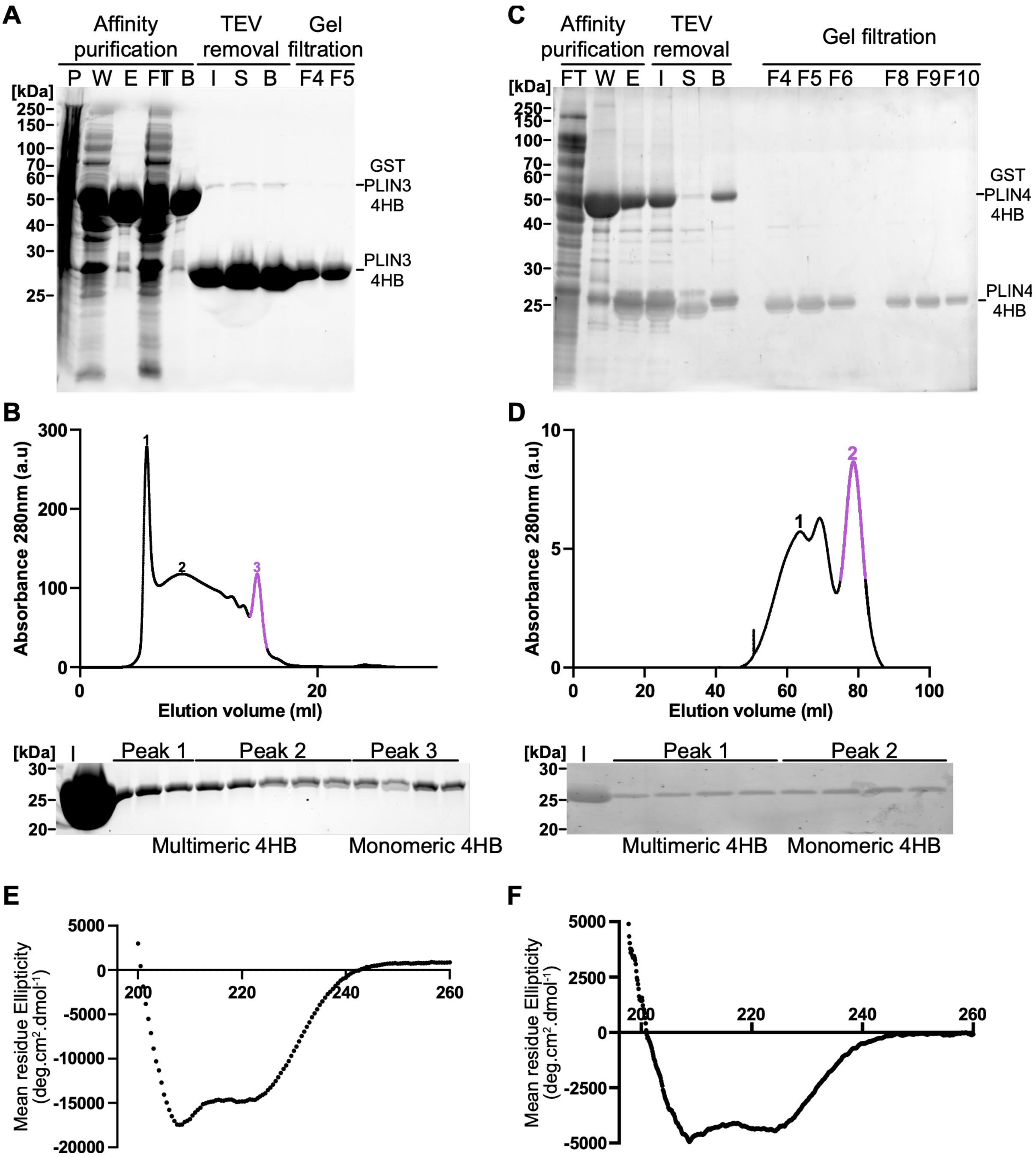


**Figure S2. Purification and analysis of PLIN3-4HB and PLIN4-4HB.**

**(A-D)** PLIN3-4HB (aa197-434) and PLIN4-4HB (aa1160-1371) were purified from *E. coli* as GST-fusion proteins by affinity chromatography on glutathione-beads, followed by elution from the beads using TEV protease to cleave the GST tag, and a final purification of the monomeric proteins by gel filtration. SDS-PAGE analysis of fractions from the purification of PLIN3-4HB (A) and PLIN4-4HB (C), stained with Sypro Orange. Fractions are labeled as: P, bacterial pellet; W, wash; FT, flow-through from the glutathion beads; E, elution fraction; I, input before TEV removal on Ni-NTA beads; S, supernatant; B, beads; F, fractions from the gel filtration columns. Sizes of the molecular weight standards are indicated. (B, D) Gel-filtration profiles of PLIN3-4HB on Superose 6 10/300 (B) or PLIN4-4HB on Hiprep 16/60 sephacryl S-300 HR (D). The columns were calibrated with protein standards (BSA: 66 kDa and Cytochrome C: 12.4 kDa) to determine the molecular weight of elution peaks; peaks marked in pink represent monomeric fractions; higher molecular weight oligomeric peaks are shown in black. Absorbance at 280 nm (a.u.) is plotted against the elution volume (mL). Fractions were analyzed by SDS-PAGE with Sypro Orange staining. Lane I represents the input sample before gel filtration. **(E-F)** CD spectroscopy analysis of the monomeric PLIN3-4HB (E) or Plin4-4HB (F) in buffer.

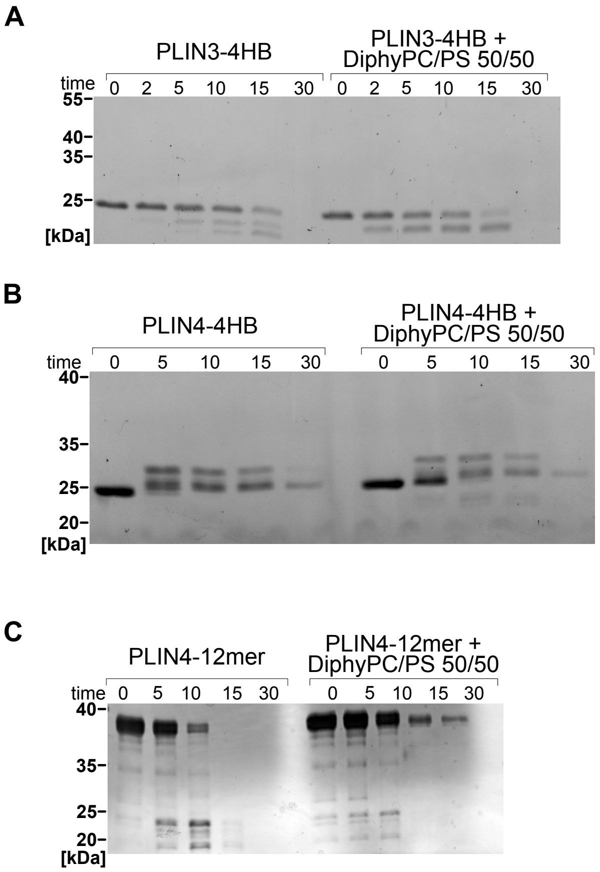


**Figure S3. Additional data related to Figure 2 (limited proteolysis).**

**(A, B)** Limited proteolysis of PLIN3-4HB (A) and PLIN4-4HB (B) in solution or in the presence of Diphy liposomes (50% PC, 50% PS) by subtilisin (0.2 μg/ml). Samples were retrieved at indicated time points: 0, 2, 5, 10, 15 and 30 min for PLIN3 and PLIN3-4HB, and 0, 5, 10 and 30 min for PLIN4-4HB. Samples were analyzed by SDS-PAGE and stained with SyproOrange. Sizes of the molecular weight standards are indicated. Gels are representative of at least two independent experiments. **(C)** Limited proteolysis of PLIN4-12mer[524] in solution or in the presence of Diphy liposomes (50% PC, 50% PS) by subtilisin (0.2 μg/ml). Samples were retrieved at indicated time points: 0, 2, 5, 10, 15 and 30 min. Samples were analyzed by SDS-PAGE and stained with silver nitrate. Sizes of the molecular weight standards are indicated. Gels are representative of at least two independent experiments.

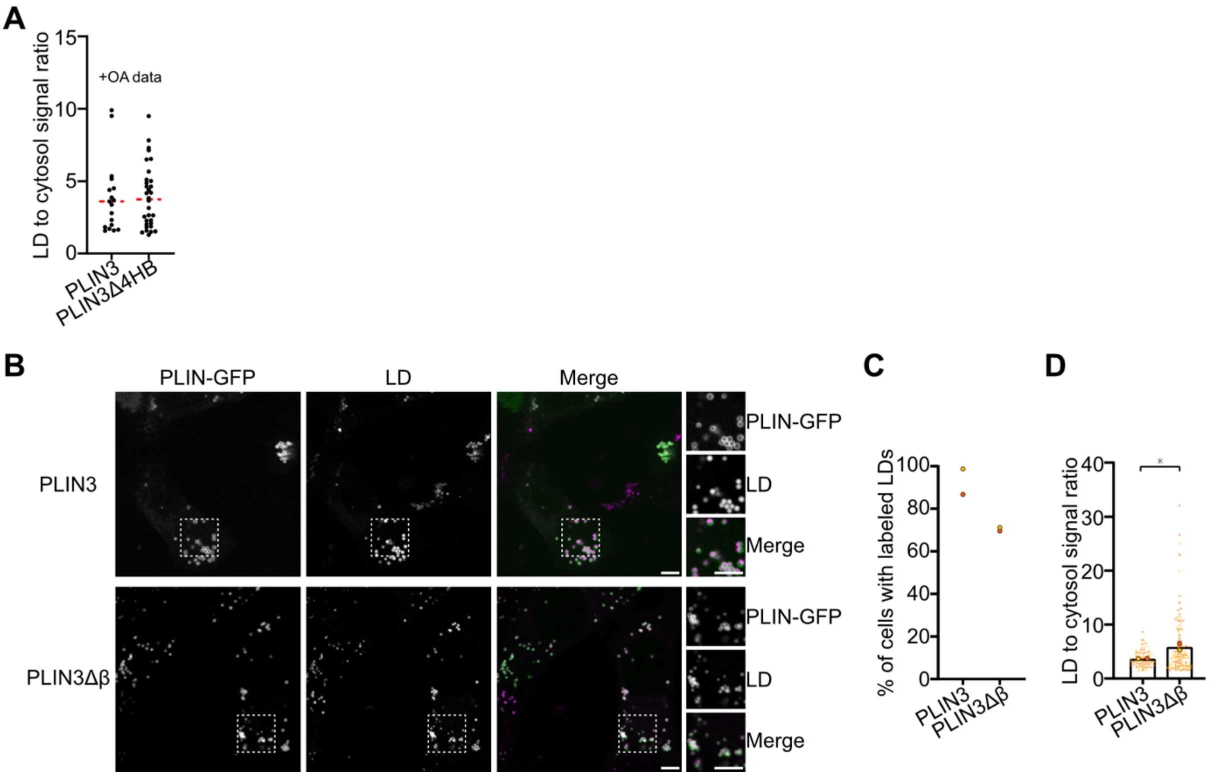


**Figure S4. Additional Data related to Figure 4.**

**(A)** Localization of PLIN3 and PLIN3Δβ, lacking the C-terminal region of the 4HB domain, upon transient expression in HeLa cells grown in standard media. Both proteins contain an N-terminal GFP tag. LDs were stained with Lipi-Blue. Magnifications (1.5x) of the indicated areas are shown on the right. Scale bars: 5 µm. **(B)** Quantification of experiments shown in (A), showing % of cells with protein signal on LDs. Data points represent independent experiments (20-30 cells per experiment), each colour representing a single experiment, lines show means or means ± SD from 2 to 5 experiments. **(C)** Quantification of experiments shown in (A), showing the ratio of GFP signal between LDs and the cytosol. Small symbols represent the mean of ratios for LDs in one cell (single z-section), large symbols represent the mean of each independent experiment, different colours are used for different experiments. Bars represent means ± SEM from 5 independent experiments. Samples were compared using a Mann-Whitney test, *P <0.05.

**Captions for Supplemental Videos**

**Video S1: Molecular Dynamics simulation of 4HB model from PLIN1 (AF).** Video shows 1 µs simulation of the 4HB model of PLIN1 generated using AlphaFold 3 (AF) server (see Methods). The structure is represented in cartoon style colored by secondary structure (alpha-helix in magenta, beta-sheet in yellow, coils in white or light-blue). The side chains are represented in lines colored by atom names. The water molecules and ions are not shown for clarity.

**Video S2: Molecular Dynamics simulation of 4HB model from PLIN1 (HM).** Video shows 1µs simulation of the 4HB model of PLIN1 generated using Homology modelling method (HM) (see Methods). The structure is represented in cartoon style colored by secondary structure (alpha-helix in magenta, beta-sheet in yellow, coils in white or light-blue). The side chains are represented in lines colored by atom names. The water molecules and ions are not shown for clarity.

**Video S3: Molecular Dynamics simulation of 4HB model from PLIN2 (AF).** Video shows 1µs simulation of the 4HB model of PLIN2 generated using AlphaFold 3 (AF) server(see Methods). The structure is represented in cartoon style colored by secondary structure (alpha-helix in magenta, beta-sheet in yellow, coils in white or light-blue). The side chains are represented in lines colored by atom names. The water molecules and ions are not shown for clarity.

**Video S4: Molecular Dynamics simulation of 4HB model from PLIN2 (HM).** Video shows 1µs simulation of the 4HB model of PLIN2 generated using Homology modelling method (HM) (see Methods). The structure is represented in cartoon style colored by secondary structure (alpha-helix in magenta, beta-sheet in yellow, coils in white or light-blue). The side chains are represented in lines colored by atom names. The water molecules and ions are not shown for clarity.

**Video S5: Molecular Dynamics simulation of 4HB model from PLIN3 (AF**). Video shows 1 µs simulation of the 4HB model of PLIN3 generated using AlphaFold 3 (AF) server (see Methods). The structure is represented in cartoon style colored by secondary structure (alpha-helix in magenta, beta-sheet in yellow, coils in white or light-blue). The side chains are represented in lines colored by atom names. The water molecules and ions are not shown for clarity.

**Video S6: Molecular Dynamics simulation of 4HB model deom PLIN3 (HM).** Video shows 1µs simulation of the 4HB model of PLIN3 generated using Homology modelling method (HM) (see Methods). The structure is represented in cartoon style colored by secondary structure (alpha-helix in magenta, beta-sheet in yellow, coils in white or light-blue). The side chains are represented in lines colored by atom names. The waters molecules and ions are not shown for clarity.

**Video S7: Molecular Dynamics simulation of 4HB model from PLIN4 (AF).** Video shows 1 µs simulation of the 4HB model of PLIN4 generated using AlphaFold 3 (AF) server (see Methods). The structure is represented in cartoon style colored by secondary structure (alpha-helix in magenta, beta-sheet in yellow, coils in white or light-blue). The side chains are represented in lines colored by atom names. The water molecules and ions are not shown for clarity.

**Video S8: Molecular Dynamics simulation of 4HB model from PLIN4 (HM).** Video shows 1µs simulation of the 4HB model of PLIN4 generated using Homology modelling method (HM) (see Methods). The structure is represented in cartoon style colored by secondary structure (alpha-helix in magenta, beta-sheet in yellow, coils in white or light-blue). The side chains are represented in lines colored by atom names. The water molecules and ions are not shown for clarity.

**Video S9: Molecular Dynamics simulation of 4HB model from PLIN5 (AF).** Video shows 1 µs simulation of the 4HB model of PLIN5 generated using AlphaFold 3 (AF) server (see Methods). The structure is represented in cartoon style colored by secondary structure (alpha-helix in magenta, beta-sheet in yellow, coils in white or light-blue). The side chains are represented in lines colored by atom names. The water molecules and ions are not shown for clarity.

**Video S10: Molecular Dynamics simulation of PLIN3-4HB$\Delta\alpha\beta$ (HM).** Video shows 1µs simulation of PLIN3-4HB$\Delta\alpha\beta$ generated using Homology modelling method (HM) (see Methods). The structure is represented in cartoon style colored by secondary structure (alpha-helix in magenta, beta-sheet in yellow, coils in white or light-blue). The water molecules and ions are not shown for clarity.

**Video S11: Molecular Dynamics simulation of PLIN3-4HB$\Delta\alpha\beta$ (AF).** Video shows 1µs simulation of PLIN3-4HB$\Delta\alpha\beta$ generated using AlphaFold 3 (AF) server (see Methods). The structure is represented in cartoon style colored by secondary structure (alpha-helix in magenta, beta-sheet in yellow, coils in white or light-blue). The water molecules and ions are not shown for clarity.

**Video S12: Molecular Dynamics simulation of PLIN3-4HB on an LD surface (R1).** Video shows 500 ns simulation (repeat 1) of PLIN3-4HB placed in water above the LD surface at the start of simulation (see Methods). The structure is represented in cartoon style. The four helices of the 4HB are colored in red (H1), blue (H2), green (H3) and yellow (H4). The extra-helix in PLIN3-4HB named H5 is colored in orange. Triolein (TG(18:1/18:1/18:1)) is colored in tan and PC(16:0/18:1) molecules in stick style are colored in blue. The blue line represents the system with periodic boundary condition. Water molecules and ions are not shown for clarity.

**Video S13: Molecular Dynamics simulation of PLIN3-4HB on an LD surface (R2).** Video shows 500 ns simulation (repeat 2) of PLIN3-4HB placed in water above the LD surface at the start of simulation (see Methods). The structure is represented in cartoon style. The four helices of the 4HB are colored in red (H1), blue (H2), green (H3) and yellow (H4). The extra-helix in PLIN3-4HB named H5 is colored in orange. Triolein (TG(18:1/18:1/18:1)) is colored in tan and PC(16:0/18:1) molecules in stick style are colored in blue. The blue line represents the system with periodic boundary condition. Water molecules and ions are not shown for clarity.

**Video S14: Molecular Dynamics simulation of PLIN3-4HB on an LD surface (R3).** Video shows 500 ns simulation (repeat 3) of PLIN3-4HB placed in water above the LD surface at the start of simulation (see Methods). The structure is represented in cartoon style. The four helices of the 4HB are colored in red (H1), blue (H2), green (H3) and yellow (H4). The extra-helix in PLIN3-4HB named H5 is colored in orange. Triolein (TG(18:1/18:1/18:1)) is colored in tan and PC(16:0/18:1) molecules in stick style are colored in blue. The blue line represents the system with periodic boundary condition. Water molecules and ions are not shown for clarity.

**Video S15: Molecular Dynamics simulation of PLIN4-4HB on an LD surface (R1).** Video shows 500 ns simulation (repeat 1) of PLIN4-4HB placed in water above the LD surface at the start of simulation (see methods). The structure is represented in cartoon style. The four helices of the 4HB are colored in red (H1), blue (H2), green (H3) and yellow (H4). Triolein (TG(18:1/18:1/18:1)) is colored in tan and PC(16:0/18:1) molecules in stick style are colored in blue. The blue line represents the system with periodic boundary condition. Water molecules and ions are not shown for clarity.

**Video S16: Molecular Dynamics simulation of PLIN4-4HB on an LD surface (R2).** Video shows 500 ns simulation (repeat 2) of PLIN4-4HB placed in water above the LD surface at the start of simulation (see methods). The structure is represented in cartoon style. The four helices of the 4HB are colored in red (H1), blue (H2), green (H3) and yellow (H4). Triolein (TG(18:1/18:1/18:1)) is colored in tan and PC(16:0/18:1) molecules in stick style are colored in blue. The blue line represents the system with periodic boundary condition. Water molecules and ions are not shown for clarity.

**Video S17: Molecular Dynamics simulation of PLIN4-4HB on an LD surface (R3).** Video shows 500 ns simulation (repeat 3) of PLIN4-4HB placed in water above the LD surface at the start of simulation (see methods). The structure is represented in cartoon style. The four helices of the 4HB are colored in red (H1), blue (H2), green (H3) and yellow (H4). Triolein (TG(18:1/18:1/18:1)) is colored in tan and PC(16:0/18:1) molecules in stick style are colored in blue. The blue line represents the system with periodic boundary condition. Water molecules and ions are not shown for clarity. Note that in this simulation, PLIN4-4HB became trapped between two boundary conditions due to the small size of the system. Because of this artefact, this simulation was excluded from further analysis.
